# A Multi-State Structural Genomics Approach Enables Large-Scale, Mechanistic, and Context-Specific Classification of ABCC6 Genetic Variants Implicated in Calcification Diseases

**DOI:** 10.1101/2025.06.17.660194

**Authors:** Jessica B. Wagenknecht, Neshatul Haque, Salomao D. Jorge, Brian D. Ratnasinghe, Raul Urrutia, William A. Gahl, Shira G. Ziegler, Michael T. Zimmermann

**Affiliations:** Computational Structural Genomics Unit, Linda T. and John A. Mellowes Center for Genomic Sciences and Precision Medicine, Medical College of Wisconsin, Milwaukee, WI 53226, USA; Linda T. and John A. Mellowes Center for Genomic Sciences and Precision Medicine, Medical College of Wisconsin, Milwaukee, WI 53226, USA; Department of Biochemistry, Medical College of Wisconsin, Milwaukee, WI 53226, USA; Department of Surgery, Medical College of Wisconsin, 8701 Watertown Plank Road, Milwaukee, WI 53226, USA; Medical Genetics Branch, National Human Genome Research Institute, National Institutes of Health, Bethesda, MD 20892, USA; Department of Genetic Medicine, Johns Hopkins University School of Medicine, Baltimore, Maryland 21205, USA; Data Science Institute, Medical College of Wisconsin, Milwaukee, WI 53226, USA

**Keywords:** Precision Medicine, Genomic Interpretation, Variant Prioritization, ABCC6, Pseudoxanthoma elasticum, Generalized arterial calcification of infancy

## Abstract

**Purpose:** Genetic variation in ATP Binding Cassette Subfamily C Member 6 (ABCC6) can cause both pseudoxanthoma elasticum (PXE) and generalized arterial calcification of infancy (GACI). Despite both diseases being rare, there are already 930 distinct missense variants in ABCC6 reported, 87% of which are of uncertain clinical significance (VUS). New approaches are needed to interpret and classify these VUS mechanistically.

**Methods:** We developed 3D protein models of ABCC6 in three functionally relevant conformations to calculate the structural effects of variants and identify 3D mutational hotspots. With this and additional functional information, we categorized variants in a mechanistic ontology based on which critical functions of ABCC6 they impact. We then compared PXE and GACI -associated variants.

**Results:** We identified two three-dimensional hotspots of pathogenic variants and six specific functions of ABCC6 which variants impact. From this, we propose a mechanism for pathogenicity for 41% of VUS according to their impacted function, 30 of which could be reclassified as Likely Pathogenic from our non-clinical data. Finally, we found slight differences between PXE and GACI-associated variants.

**Conclusion:** The mechanistic information we present will guide future research to better address calcification disorders and understand genetic variants. Further, our VUS reclassification will improve the diagnosis of ABCC6-driven diseases, shortening diagnostic odysseys. We believe that computational structural genomics approaches will soon take prominence in genomics data interpretation.

**Highlights:** Variant 3D hotspot detection and effect clustering, GACI and PXE Variant Comparison, consistent ability to (re)categorize known pathogenic variants, and population genetics differences.

## Introduction

ATP-Binding Cassette (ABC) proteins are among the most ubiquitous transport proteins, harnessing ATP hydrolysis to translocate substrates across vesicle and plasma membranes. While proteins in ABC subfamily C are most known for their role in drug resistance, they can also cause heritable diseases when mutated^1^. Loss-of-function variants of the *ABCC6* gene (HGNC:57) have been associated with the recessive disease pseudoxanthoma elasticum (PXE; OMIM #264800 and #177850) and, less frequently, Generalized Arterial Calcification of Infancy (GACI; OMIM #614473). Both diseases are characterized by ectopic calcification. In PXE, progressive calcification of the elastic fibers in the skin, eyes, and arteries cause dermal papules and angioid streaks, although PXE is typically not lethal. On the other hand, patients with GACI have extensive vascular calcification that leads to 50% mortality by six months of age^2, 3^. Notably, GACI is more frequently caused by Ectonucleotide Pyrophosphatase/Phosphodiesterase 1 (ENPP1) deficiency. While genetic variation in *ABCC6* has long been understood to underly PXE and, more recently, GACI^4^, the molecular mechanisms affected by its dysregulation are not fully understood. This is partly because the normal physiologic function of ABCC6 remains elusive despite recent work studying the role of ABCC6 in ATP and nucleotide transport^5–7^. This lack of a precise mechanism adds to the challenge of diagnosing and treating PXE and GACI. Moreover, there are hundreds of rare variants in *ABCC6* whose clinical significance is uncertain (VUS; n = 782)^8^; far fewer are clinically classified as benign (n = 41) or pathogenic (n = 84). The large number of VUS needing interpretation leaves many individuals without clear answers about their health status, limiting diagnostic and therapeutic research options.

Our work to increase the scale and rigor of computational structural genomics^9–12^, and increased adoption of similar approaches by other labs^13, 14^, demonstrates the high value of a proteogenomic approach that leverages structural bioinformatics to interpret the mechanisms of dysfunction of missense genetic variation. We anticipate that the ever expanding national and international genomics datasets will continue to increase the need for more accurate approaches to mechanistically interpreting human genetic variation.

While there is currently no full-length experimental structure of ABCC6, information from other members of its conserved family of transport proteins can be leveraged to predict the structure of ABCC6 and infer basic features of its catalytic and transport mechanisms. ABCC6 is a long-type multidrug resistance protein^15^, meaning it contains two transmembrane domains (TMD1 and TMD2), two nucleotide-binding domains (NBD1 and NBD2), and an initial transmembrane domain (TMD0)^16^. TMD0 is linked to TMD1 through a loop designated L0; a second loop, L1, links NBD1 to TMD2 (**Figure 1A**). TMDs 1 and 2 create a gated channel through the cellular membrane, open to the intracellular side at baseline, allowing ligand binding. When two ATP-Mg^+2^ bind at the interface between the two cytoplasmic NBDs, the extracellular facing conformation is formed, completing transport. ATP hydrolysis returns the protein to its initial state (**Figure 1B**). Thus, despite the lack of experimentally solved structures, there is considerable opportunity to apply structural bioinformatics approaches to mechanistically interpret ABCC6 genetic variation.

**Figure 1:**
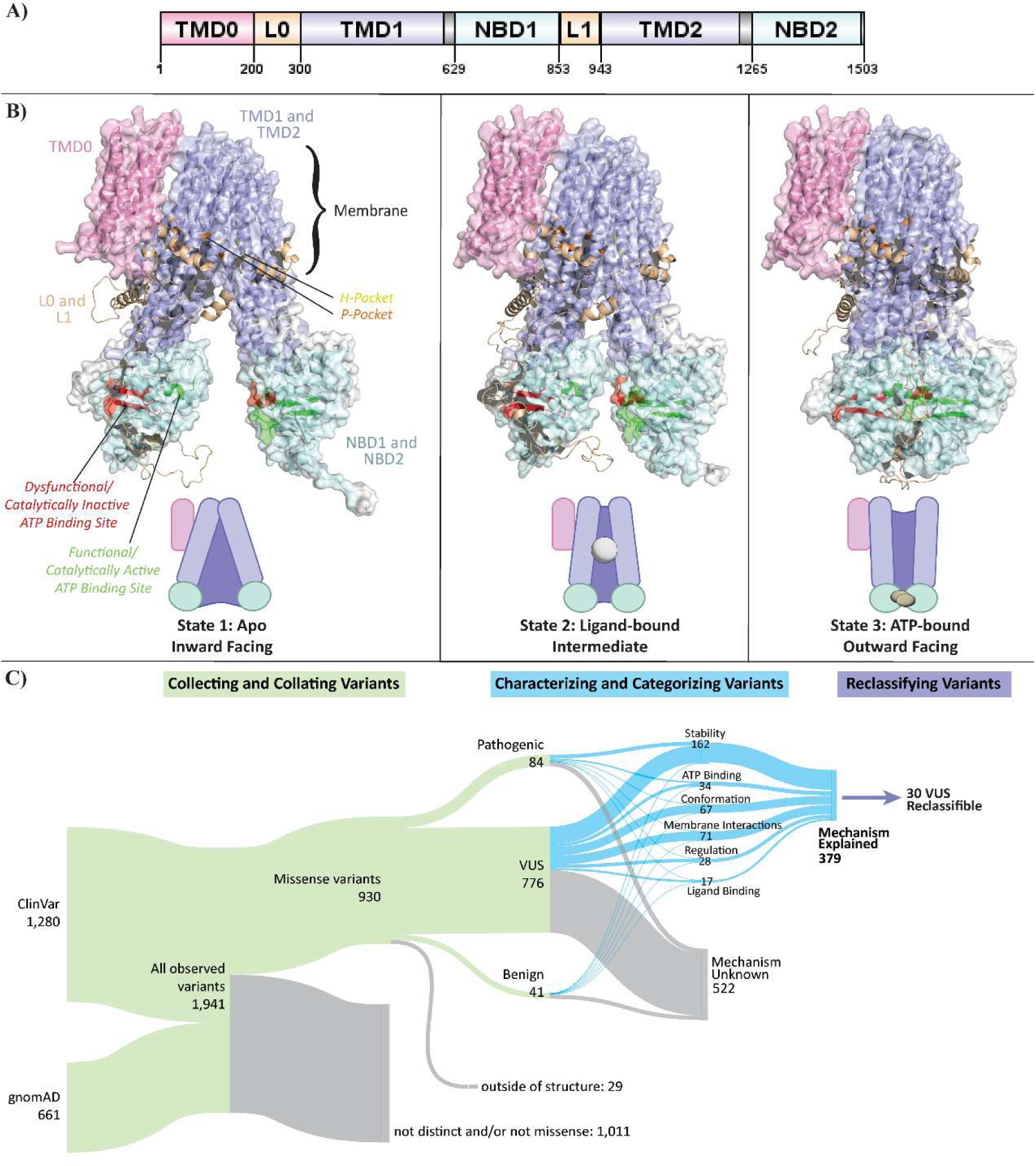
The interpretation and classification of genetic variants in ABCC6 is greatly enhanced by our novel process, which uses 3 state models of the protein. A) The domains of ABCC6 are shown linearly along the protein sequence where Transmembrane domains 1 and 2 (TMD1 and TMD2) are colored purple, Nucleotide binding domains 1 and 2 (NBD1 and NBD2) are light blue, Loop 0 and 1 (L0 and L1) are colored tan, and Transmembrane domain 0 (TMD0) is pink.B) ABCC6 was represented in three states: on the left, the inward-facing and apo conformation (also called “state 1”); in the middle, the intermediate or ligand-bound conformation (also called “state 2”); and on the right, the outward-facing or ATP-bound conformation (also called “state 3”). In the diagram, the domains are colored the same as in B. Further, the P-pocket is colored orange, the H-pocket yellow, the catalytically active nucleotide binding site is colored green and the inactive site red^17, 18^. C) The flow of variants through our process, as described in Methods, is shown as a Sankey plot. Variants are first collected and collated to those within our structures, then characterized and categorized by their structural and functional information, which was used to identify the mechanism by which variants impact ABCC6 function. This could be used as evidence for ACMG variant interpretation lines PM1 and PP3 to reclassify at least 30 variants from our non-clinical data alone.

The current study uses a proteogenomic approach, leveraging multidisciplinary, mechanical, and functional information to characterize 901 ABCC6 missense genetic variants. Our approach includes calculating state-specific scores to harness the dynamic nature of the transport process and identifying all key functional regions. From these investigations, we categorize 319 (41%) VUS by their likelihood to alter specific protein features. Additionally, we propose that 30 VUS can be reclassified as likely pathogenic according to their underlying molecular mechanisms. Another 134 VUS, with the additional evidence we propose, are on the cusp of a likely pathogenic interpretation. Through our process, we aim to better understand which variants may cause PXE and GACI. Finally, we anticipate that by accounting for molecular context, such as we have herein through integrative 3D modeling and biophysical characterizations, more genetic variants can be clearly categorized, functional genomics tests better designed, and clinical diagnostics improved.

## Materials and Methods

### Integrative Three-State Protein Structure Modeling Captures Details of ABCC6 Function

There are no full-length experimentally determined models of ABCC6. We used integrative modeling to generate the first models of ABCC6 for use in scoring genomic variation. ABCC6’s closest homologous protein, ABCC1, has a series of 3 bovine models published in the Protein Data Bank (PDB)^17–19^ that show the range of motion ABCC1 goes through at the un-liganded inward-facing conformation to an intermediate conformation with substrate bound, and then to the outward-facing conformation with ATP bound at the hydrolytic domain. After these three steps, ABCC1 hydrolyzes ATP to return to its apo conformation. Because each structure highlights an essential step in ABCC1’s function, and ABCC6 is thought to have a similar series and mechanism of motion, we used each structure as a template in Modeller^20^ to generate 3D models of ABCC6 in 3 functional states. However, the ABCC1 models (and consequently ABCC6 models) lacked the TMD0 domain of ABCC6. YCF1, the yeast ortholog of ABCC1, has two structures on the PDB^21, 22^, which include a well-resolved TMD0 and have better resolution on L0 and L1 than ABCC1’s structures. Therefore, these two structures were used as templates with Phyre^23^ to create two models of ABCC6 in the inward-facing conformation (state 1) and intermediary conformation (state 2). There is no structure of outward-facing (state 3) YCF1, but TMD0 and L0 had negligible conformational differences between states 1 and 2, so an improved series of models were created by replacing the TMD0 and L0 domains from the ABCC1-template structures with the YCF1-template state one domains, and replacing L1 in states 1 and 2 with the YCF1-template model L1 (but retaining the ABCC1-template L1 in state 3) (**Figure 1B**). PyMOL was used to visualize structures^24^. These steps combined generated molecular models for mechanistic interpretation of ABCC6 genetic variations.

### Genomic Variation Collection and Annotation

We generated three datasets of genomic variation. First, our team selected ten missense variants from our clinical cohort with deep clinical phenotyping data. Second, we assembled 1280 ABCC6 variants from the May 2023 ClinVar release^8^ and 661 missense ABCC6 variants from gnomAD v3.1.222 using the canonical transcript (ENST00000205557). Nomenclature was validated with VariantValidator^25^. Annotations about each variant, such as associated phenotype, pathogenicity, and allele frequency, were provided by ClinVar and gnomAD, respectively (when available). Ensembl’s^26^ Variant Recorder was used to obtain variant HGVS Protein labels for ClinVar variants. These data were collated into one dataset of 930 distinct missense variants to compare information and context on each variant. We divided this dataset into two subsets: 1) all 84 pathogenic (and likely pathogenic) missense variants from ClinVar and 2) the 782 missense variants that are annotated as variants of uncertain significance (VUS) or those for which there were no clinical annotations available. However, six variants were at amino acids not shown in our structures (which began at residue 26). These variants were removed, leaving 84 pathogenic missense variants and 776 VUS, which we analyzed further (901 total variants). Additionally, our data included 41 (likely)benign variants. Finally, the 23 variants with conflicting interpretations of pathogenicity were not analyzed.

### Annotating Motifs, Biochemical Regions, and Post Translational Modification Sites

We first gathered sequence-based information about ABCC6’s motifs and post-translational modifications (PTM) to understand the mechanistic effects of the above variants. First, within the nucleotide-binding domain, there are five highly conserved motifs across the ABC family: Walker A (also called P-loop or phosphate-binding loop), Walker B, Switch region (also called H-loop or histidine loop), signature motif (also called C-loop), and Q-loop. These motifs, among other residues, form the nucleotide-binding pocket, which classically consists of the Walker A and B, Q-loop, and H-loop from one NBD and the signature motif of the other NBD^27–29^. Variants within these motifs, or spatially supporting each motif, could affect motif functions. This has been documented already, as ABC Subfamily C (ABCC) proteins all carry a mutation of Glu778 to Asp in the Walker B sequence of NBD1, causing decreased efficiency of ATP hydrolysis for that ATP binding site ^17, 30, 31^; hence, NBD1 is considered dysfunctional. Similarly, ABCC proteins are missing 10-13 highly conserved residues in NBD1, residues which in other ABC proteins stabilize the TMD-NBD interface; due to this loss of essential contacts, ABCC proteins are less stable in this region^18, 31^. Therefore, not only were the residues within TMD-NBD interface binding regions and Walker motifs noted, but those within the dysfunctional NBD1 and NBD1-TMD interface were also noted as being especially sensitive to variation.

Linker regions are also regulatory domains. Within L1, also sometimes called the regulatory domain or R-domain, lies a highly conserved phosphorylation site at S902, where studies in YCF1 (a yeast homolog for ABCC1) demonstrated PKA phosphorylation of that site to regulate YCF1 activity by inducing structural changes upon phosphorylation. L1 also contacts NBD1 through Q906 binding with Q715, stabilizing and regulating activity^21^. Beyond the PTMs in L1, residues 911-931 are also known to be of regulatory importance ^32^. Within L0, a CK2 phosphorylation site is highly conserved at S244, which negatively regulates substrate transport, and a predicted basolateral localization sequence aids in proper localization ^21, 33^. All these residues, as well as residues within three amino acids of a PTM, were considered of regulatory importance. Finally, there are 13 other PTMs without an experimentally-discovered function suggested by both iPMTnet^34^ and PhosphoSitePlus^35^, 9 of which are phosphorylations, two acetylations, one ubiquitination (on the same amino acid as one of the acetylations), and one n-glycosylation. Any variants sequentially or spatially near these PTMs were noted as well.

### Structure-Based Biophysical Annotations and Fold Stability

Since the ABCC1 substrate and ATP binding pockets are well-defined^17, 18, 30, 31, 36^, sequence alignment was used to predict the ABCC6 substrate binding sites. To obtain further 3D contacts for both ATP and the ligand, we aligned the template ABCC1 structures with our ABCC6 structure in states 2 and 3 to confirm the homology-based prediction of ligand-binding pockets. We defined the pockets as the union of conserved ligand-binding sites, ATP-binding sequence motifs, and all residues within 5 Å of aligned ABCC1 experimentally-determined ligand positions.

The membrane-spanning portion of ABCC6 was predicted with the PPM 3.0 webserver^37^ and all residues within this region and an accessible surface area of > 50% were tagged as lipid-interacting. We also noted three positively charged helices, which rest upon the membrane’s outer layer at residues 204-208, 217-228, and 931-943. These residues were tagged for their likely role in charge-stabilizing membrane head group interactions.

FoldX^38^ was used to obtain the change in folding free energy (ΔΔG_fold_) for each variant compared to the wildtype structure. The same procedure was performed with all three conformations as input and across all pathogenic variants from all three cohorts. For the 10 cohort variants, FrustratometeR^39^ was also used to obtain the change in folding frustration energy. Finally, for all variants, the PAM30 score for the replacement was obtained, as well as the difference in sidechain hydrophobicity using the Peptides R package^40^ and the Kyte-Doolittle scale^41^.

### Discovery of 3D Genetic Variant Hotspots

We determined amino acid contact maps using those within 10 Å of each amino acid in the state 1 and state 3 structures. Since 39 of the 84 pathogenic variants are also reported in gnomAD, we performed two rounds of hotspot analysis. First, we weighted variants by their population incidence by summing the allele counts for all gnomAD-reported variants within 10 Å of each residue. Second, we weighted distinct variants by summing the total number of variants (out of the 84 total pathogenic variants) within 10 Å of each residue. The residues with 17 or more alleles or six or more variants, respectively, were considered very variant-rich, and those with 12 or more alleles or five variants were considered fairly variant-rich. To calculate statistical significance, we first made two null sets with the same number of randomly selected variants as each of our observations (total allele count of 269 and 84 total variants, respectively) and repeated a similar procedure, permuting selection 10,000 times to obtain the number of times that randomly distributed variants were more densely populated around each residue than our observations. Dividing that number by 10,000 gave an empirical p-value.

### Categorizing Mechanistic Effect of Variants

To characterize the range of effects that genetic variants have on ABCC6, we describe six essential aspects of ABCC6 function: ATP Binding, Ligand Binding, Membrane Interactions, Regulation, Conformational Dynamics, and Functional Stability (see Table 1 for an explanation of categories). These six categories of functional effect were chosen based on our study of wild-type ABCC6 and a cursory review of the variants’ impacts. They formed the basis of our ABCC6-specific context-guided variant effect algorithm – a unique approach to variant effect interpretation that focuses on understanding the mechanism of variant impact rather than correlating features to clinical information. This decision-tree algorithm uses all the annotations and scores described in prior sections to categorize variants into one of the six categories (see Supplementary Materials for the code). Its rules are logically derived: a series of metrics were chosen to define each function, and thresholds set for those metrics. Then, each function is tested based on order of specificity – for example, if a variant is destabilzing, and destabilizing to the ATP binding region, it’s impacting ATP binding specifically. We tested our algorithm on the cohort variants and compared it with the manual categorization we performed, then applied it to missense pathogenic variants, benign variants, and VUS. R packages ComplexHeatmap^42^ and Tidyverse^43^ were used in our analysis. Finally, other pathogenicity prediction scores from CADD^44^, REVEL^45^, PolyPhen2^46^, AlphaMissense^47^, and SIFT^48^ were obtained through their respective servers or databases. The threshold for which variants were predicted to be pathogenic was set at their respective recommended thresholds: ≥ 20 for CADD (using the PHRED score)^44^, ≥ 0.5 for REVEL^45^ and PolyPhen2^49^, ≥ 0.564 for AlphaMissense^47^, and ≤ 0.05 for SIFT ^49^.

**Table 1:** Our categorization process focuses on delineating the mechanistic impacts of a variant on ABCC6, while the ACMG classification of a variant describes its likelihood of causing a specific disease. Therefore, each variant was given both a category of mechanistic impact (if possible), as well as a class of pathogenicity. Furthermore, the categorization of a variant was used as PP3 criteria of pathogenicity.

| Methodology | Groups/Terms | Definition/Meaning |
| --- | --- | --- |
| Categorizing variants via our protein-based structural genomics method.<br>Categorizes variants by mechanistic impact. | ATP Binding | Variants impacting the ability of ATP to bind in the nucleotide binding domains |
|  | Ligand Binding | Variants impacting the ability of the ligand to bind in the conserved binding domain |
|  | Membrane Interactions | Variants impairing the stability of ABCC6 within the basolateral membrane |
|  | Regulation | Variants impairing mechanisms of regulating ABCC6 expression and activity |
|  | Conformational Dynamics | Variants hindering ABCC6 from moving between key conformations |
|  | Functional Stability | Variants destabilizing ABCC6 overall |
| Classifying variants via the ACMG guidelines for interpretation of variant effect.<br>Classifies variants by likelihood to cause disease. | Pathogenic | Variants considered disease-causing for a specific disease |
|  | Likely Pathogenic | Variants with >90% certainty of being disease-causing for a specific disease |
|  | Variant of Uncertain Significance (VUS) | Variants which may or may not cause a specific disease |
|  | Likely Benign | Variants with >90% certainty of not causing a specific disease |
|  | Benign | Variants not considered to cause a specific disease |

### Reclassifying Variants of Uncertain Significance

The American College of Medical Genetics and Genomics (ACMG) guidelines for interpreting variant effects were used to organize evidence for VUS classification ^50^. We propose that our computational structural genomics approach can provide evidence types PM1 and PP3. How to apply computational structural genomics to these additional lines of evidence for clinical interpretation is a crucial advancement from the current study. We directly compare our categorization of variants to the existing classification (**Figure 1C** and **Table 1**). For additional comparisons to our novel variant classification approaches, we also used evidence lines PM2, PM5, PP2, and PP4, established in the field and obtainable through publicly available data. For PM1, variants were noted that reside within the well-defined Walker A motif, Walker B motif, Signature Loop, H-loop, and Q-loop^27, 28, 30^ as well as variants residing within the variant-rich three 3D-hotspots identified herein. For PP3, all variants predicted by our methodology to impact key ABCC6 functionalities were noted. For PM2, variants that were present within the ClinVar database but not gnomAD were noted^8, 51^, as well as those with an allele frequency less than 0.33% and at residues past 250. The allele frequency was chosen as 0.33% since the carrier frequency for PXE is expected to be between 1 in 150 and 1 in 300 people ^52^, and the carrier frequency for GACI is expected to be around 1 in 200 people^53^. Residues before 250 were not included since the coverage of residues 0-250 in gnomAD was lacking, i.e., the fraction of individuals with coverage over 30 was less than 50%. The coverage of ABCC6 within gnomAD v3 was confirmed to be of sufficient read depth. For PM5, all variants at the same residue position as a pathogenic variant from the ClinVar database were noted. Finally, for PP4, all variants with an associated phenotype of PXE, GACI, or both in ClinVar were noted.

## Results

We used a proteogenomic approach^56^, applying structural bioinformatics and computational biochemistry to analyze ABCC6 missense genomic variation. The full dataset contains 84 pathogenic variants from ClinVar, 776 VUS from the union of ClinVar and gnomAD, and 41 benign variants from ClinVar. Our study was initiated from deep structural analysis of the ten variants from our patient cohort, eight of which are pathogenic and two are VUS.

### Categorizing Clinical Genetic Variants into Three Mechanisms of Dysfunction

Reviewing the variants identified in our clinical cohort revealed three main mechanisms through which genetic variants affected the protein: 1) preventing conformational changes; 2) directly destabilizing the protein structure; and 3) disrupting ATP binding (**Table 2**). Variants impair conformational dynamics by changing the transmembrane domain stability in one state, impacting its ability to move into the next state. For example, replacing the glycine with a serine at position 1042 creates new interactions between adjacent helices in state 3, impairing ABCC6’s ability to reset to its state 1 conformation after ATP hydrolysis (**Figure 2A)** (NP_001162.5:p.G1042S). On the other hand, variants that destabilize the protein structure in different regions or multiple states will not impact ABCC6’s ability to move between conformations but still damage its structural integrity and, therefore, ability to function as efficiently. For example, Arg1138 in TMD2 is necessary to stabilize the TMD-NBD joint due to arginine’s capacity to make multiple bonds with nearby residues; when arginine is replaced with glutamine (variant NP_001162.5:p.R1138Q), it loses essential bonds (such as with Glu679 in NBD1) that hold the two domains together in the proper orientation (**Figure 2B**). Finally, any nonconservative variant within 5 Å of the ATP binding site in 3D space will likely disrupt ATP binding by losing site-specific properties. One example is Gly1302, which is nearby (3.8Å) to ATP. Replacing small glycine with large and charged arginine causes many steric clashes, contorting the ATP binding site and hindering ATP binding (**Figure 2C**) (variant NP_001162.5:p.G1302R). Overall, inspection of each variant’s effects on ABCC6’s structure and function reveals mechanisms by which all ten disease-associated variants impact ABCC6 in ways we can both measure and biochemically explain.

**Figure 2:**
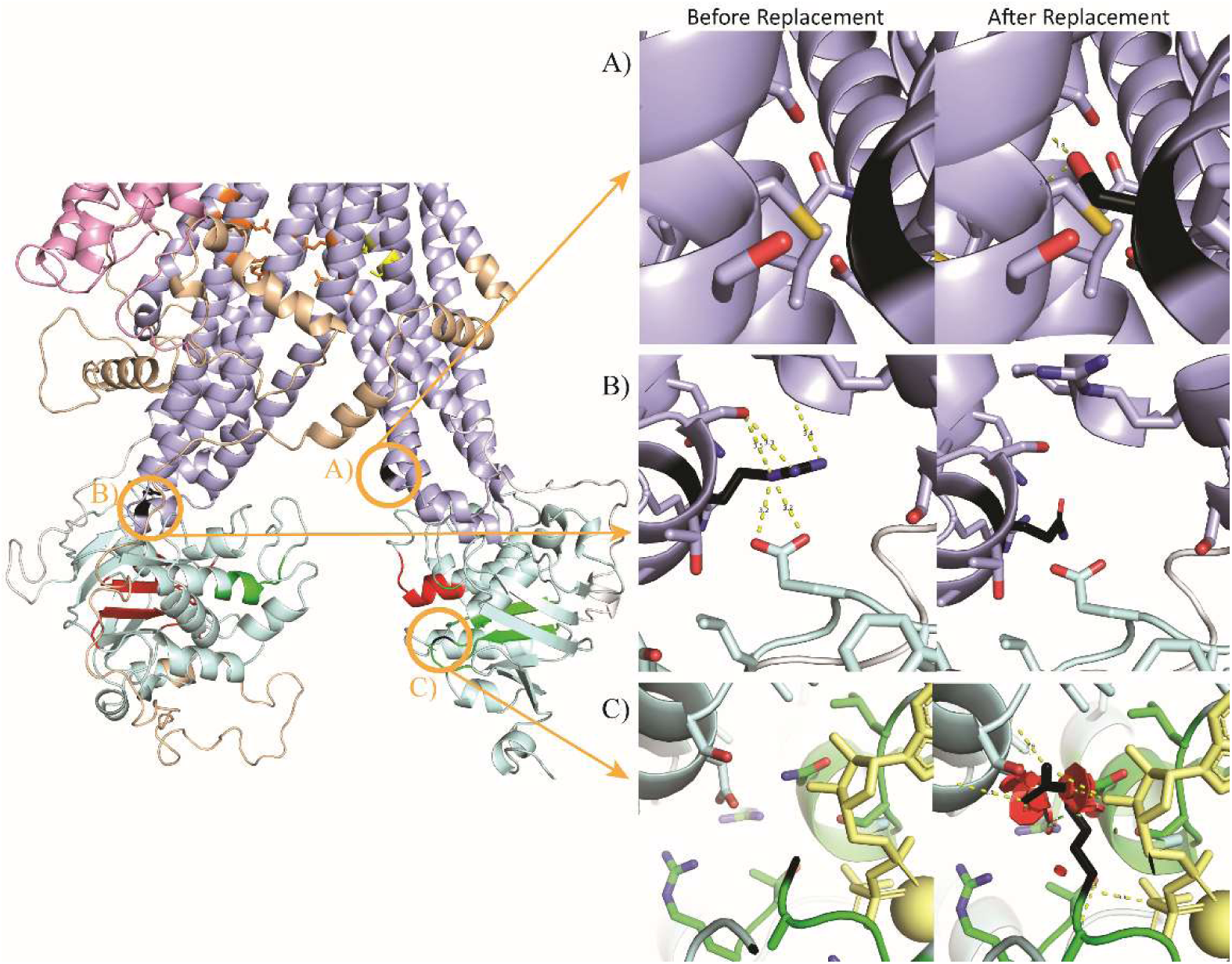
Structural modeling of ABCC6 reveals three specific mechanisms of variant effects. **A)** NP_001162.5:p.G1042S (black) disrupts conformational dynamics, as shown within the state 3 model. The serine replacement will interact with the backbone of the helix opposite to it – a helix that, despite having five serines within a 9-residue range, does not normally interact with its opposite helix. As these helices are not supposed to interact but instead be free to fall back to the state 1 conformation after hydrolysis, this substitution will inhibit the dynamics of ABCC6. **B)** NP_001162.5:p.R1138Q (black) disrupts structural stability, as shown in the state 2 model, with many bonds connecting the TMD’s arginine to the NBD’s atoms. This arginine stabilizes the crucial joint between the two domains; when this arginine is lost for an uncharged glutamine, all these bonds are lost, and the structural stability of this joint is lost in all three states. Therefore, this variant disrupts the stability of ABCC6. **C)** NP_001162.5:p.G1302R (black) disrupts ATP binding, as shown in the state 3 model. This glycine is close to many sidechains and ATP at the turn of a loop. Replacing glycine with arginine causes many steric clashes, which would cause the nucleotide-binding site to disform and the nucleotide to not bind within the pocket. This will disrupt the ATP binding of ABCC6.

**Table 2:** Single amino acid variants disrupt local conformation, ATP binding, and molecular dynamics of ABCC6. The change in energy of folding (as calculated by FoldX) and energy of conformation (as calculated by FrustratometeR) describe the impact of a variant on the stability of ABCC6, where a more positive score is more destabilizing, and a more negative score is more stabilizing.

| Protein Variant | ΔEnergy of Folding (kcal/mol) | | | ΔEnergy of Conformation (kcal/mol) | | | Allele Frequency (from gnomAD) | Clinical Significance (from ClinVar) | Domain | Predicted Affected Function |
| --- | --- | --- | --- | --- | --- | --- | --- | --- | --- | --- |
|  | State1 | State2 | State3 | State1 | State2 | State3 |  |  |  |  |
| **S317R** | 0 | 0.1 | -1.4 | -1.1 | -0.9 | -0.9 | - | Pathogenic | TMD1 | Structural Stability |
| **G755R** | -0.7 | -0.3 | -0.1 | 2.4 | 3.2 | -3.7 | $3.30 \times 10^{-5}$ | Pathogenic | NBD1 | ATP Binding |
| **G1042S** | -0.1 | 0 | 8 | 5.4 | 2.5 | 5.2 | $5.30 \times 10^{-5}$ | Uncertain Significance | TMD2 | Conformational Dynamics |
| **T1130M** | -0.7 | -1.4 | -1 | 1.2 | 1 | -0.3 | - | Pathogenic | TMD2 | Structural Stability |
| **R1138Q** | 1.2 | 0.5 | 0.4 | 1.1 | 1.1 | 0.7 | $5.90 \times 10^{-5}$ | Pathogenic | TMD2 | Structural Stability |
| **G1302R** | 1.6 | 3.3 | 3.6 | 4 | 3.9 | 10.4 | $3.90 \times 10^{-5}$ | Pathogenic | NBD2 | ATP Binding |
| **R1114C** | 1 | 1.4 | 2 | -0.6 | 1.2 | 1.8 | $3.90 \times 10^{-5}$ | Pathogenic | TMD2 | Conformational Dynamics |
| **G1263R** | 5.6 | 10.9 | 6.5 | -3.2 | -3.4 | -7.8 | $4.00 \times 10^{-5}$ | Uncertain Significance | NBD2 | Structural Stability |
| **G1296D** | 0.5 | 5.1 | 2.2 | -0.8 | 0 | 1 | $5.90 \times 10^{-5}$ | Pathogenic | NBD2 | Structural Stability |
| **R1314W** | -0.5 | -1.8 | 5.2 | -0.4 | -1.3 | 0.2 | $6.00 \times 10^{-5}$ | Pathogenic | NBD2 | Structural Stability |

Guided by the above considerations and our understanding of WT ABCC6 functioning, we developed an algorithm for systematically categorizing the mechanistic effect of ABCC6 variants. The 10 cohort variants were run through the algorithm, and eight were predicted to have the same mechanistic effect as we had assigned through our manual study. The remaining two, NP_001162.5:p.S317R and NP_001162.5:p.T1130M, were noted above as having mild differences by our manual review but were not classified by our algorithm. This provides evidence that our new algorithm can categorize ABCC6 genetic variants with precision comparable to an in-depth manual study.

### Pathogenic Variants Mostly Affect ATP Binding or Structural Stability and Cluster in Three Hotspots

We analyzed the effect of pathogenic variants on ABCC6 functionality since these are all likely to have a deleterious impact on the functioning of ABCC6. First, we analyzed whether patterns differentiate the PXE and GACI variants (**Figure 3**). Interestingly, TMD2 contains the largest portion of GACI/PXE variants (47%), while most variants associated with just PXE are in NBD1 or NBD2 (34% and 27%, respectively). NBD1 and NBD2 contain many ATP-binding residues, while NBD1 specifically has many canonical residues that destabilize ABCC proteins (compared to other ABC-family proteins). On the other hand, TMD2 contains many membrane-interacting and ligand-binding residues. However, one trend observed across both phenotypes was towards instability - almost half (46%) of the pathogenic variants’ most significant effect was destabilizing the protein, while less than 4% of the variants’ most considerable effect was stabilizing (**Figure**). Second, we predicted the likely impact of ABCC6’s pathogenic variants on its core functions, and we categorized over half (58%) with high confidence (**Figure 5**). Interestingly, more variants without a reported phenotype could be predicted than those associated with PXE or GACI. Most variants impact ABCC6’s ability to bind ATP or its structural stability – PXE variants more frequently influenced ATP binding, while variants unassociated with any phenotype more often affected structural stability.

**Figure 3:**
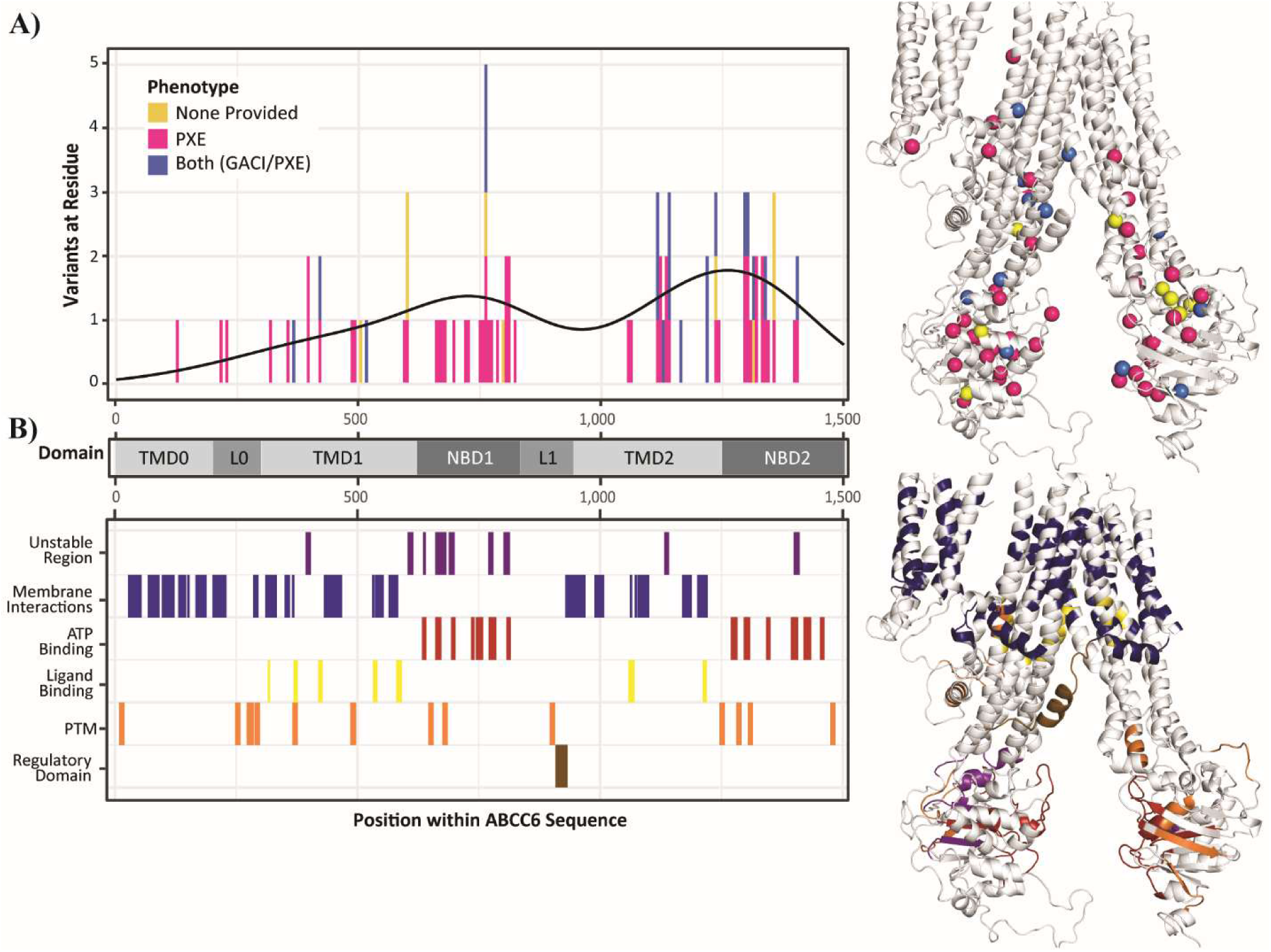
Many pathogenic variants are spatially and sequentially near functionally significant residues across ABCC6, with the majority of PXE variants found in the nucleotide-binding domains and the majority of GACI/PXE variants in the transmembrane domains. A) Demonstrates the location of ABCC6 variants across the protein (shown here in state 1), with the black line of distribution shown, while B) shows the location of functionally significant zones, including domains and regions with known functional impact. Both have the same scale graphically and the same model structurally for ease of comparison.

**Figure 4:**
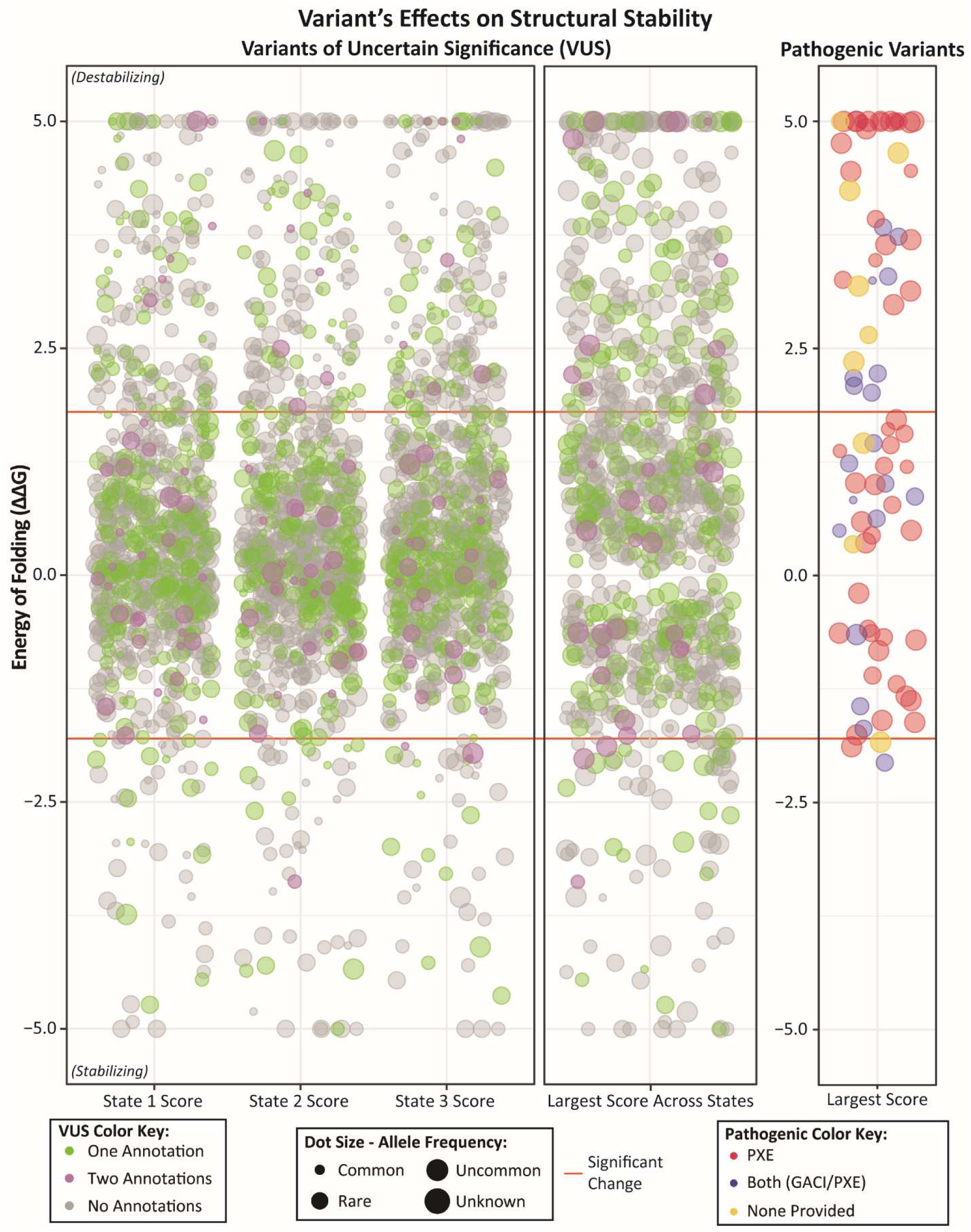
Structural Stability of ABCC6 in 3 functional states reveals that pathogenic variants and, to a lesser extent, VUS are most often destabilizing. The “largest score across states” corresponds to the highest or lowest value (whichever has the greater absolute value) for energetic stability across all three states. Hence, variants with the largest score as highly destabilizing are highly destabilizing in at least one state. Orange horizontal lines denote a significant change (in either direction) from WT, at - 1.8kcal/mol and 1.8kcal/mol.

**Figure 5:**
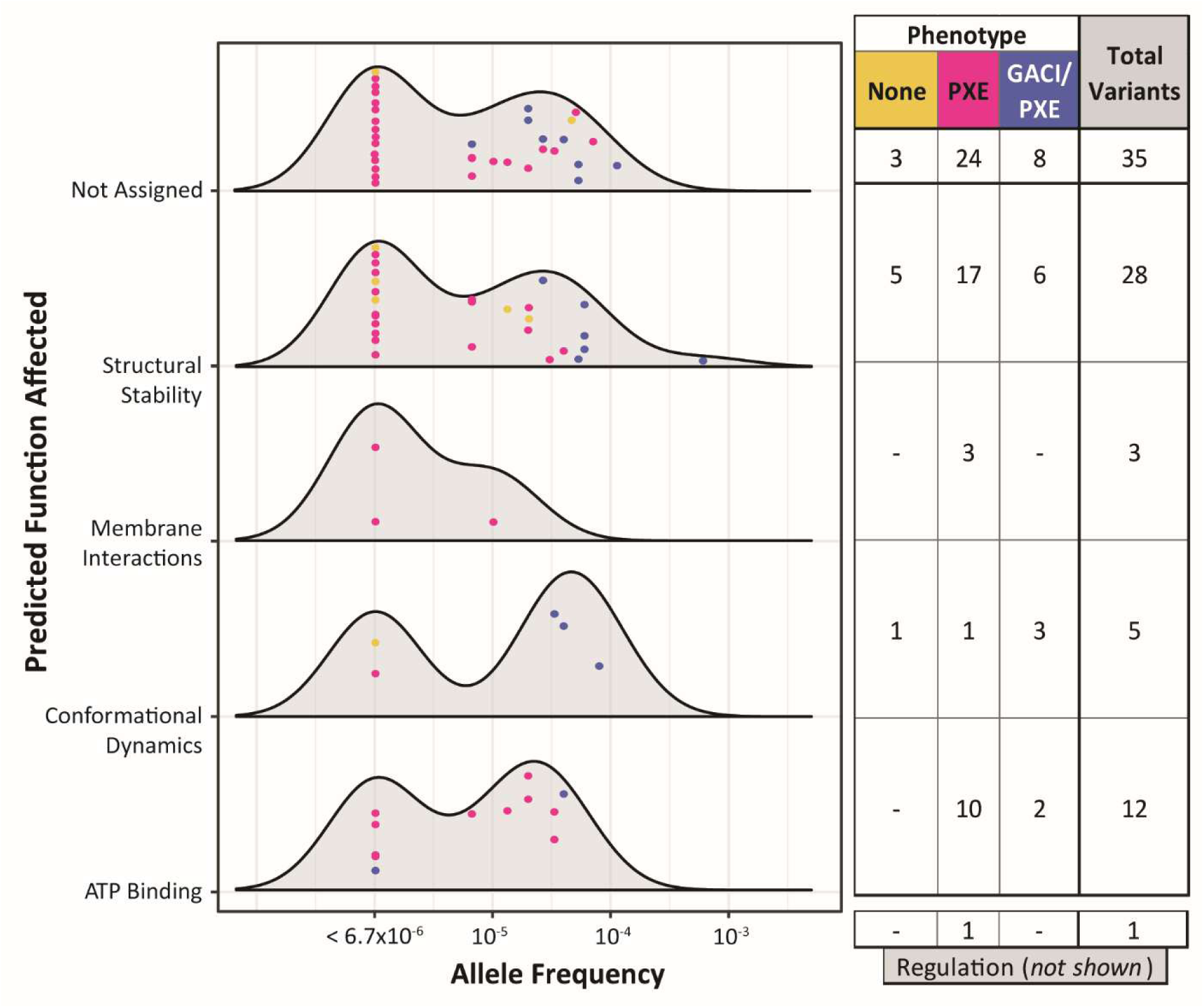
Most pathogenic variants are confidently predicted, usually affecting ATP binding or structural stability. Variant allele frequencies are shown by each structure-based function class of ABCC6 that they likely impact. Individual variants are shown within the smoothed distribution and colored by their associated phenotype from ClinVar. Total counts for variants within each effect group are shown on the right.

We performed a 3D hotspot analysis of pathogenic variants to reveal any deeper pattern across variants. Since we only had the allele count for some pathogenic variants, we counted variants in one region and the total allele count of variants in one area. Despite seeking hotspots of two different groups of variants, and in both states 1 and 3, we found hotspots in the same three regions in all analyses – one hotspot within each NBD and one spanning between the divide in the TMD bundles (**Figure 6; Supplementary Text**). These hotspots were statistically significant, with a P-value less than 0.0035 at any residues we considered within a hotspot. In fact, 30% of the top 50 residues (in terms of proximity to the most variants or variants with the highest allele count) are the same among all four analyses. It is worth noting that these three locations are incredibly functionally significant: the first two in the NBDs are where ATP binding and hydrolysis facilitate ligand movement out of the cell, and the third hotspot is in the TMDs along the path where the ligand would travel and transiently interact with ABCC6. Thus, the genetic variations diffuse along the linear gene sequence form 3D hotspots that affect the channel protein’s nucleotide and ligand binding capacities.

**Figure 6:**
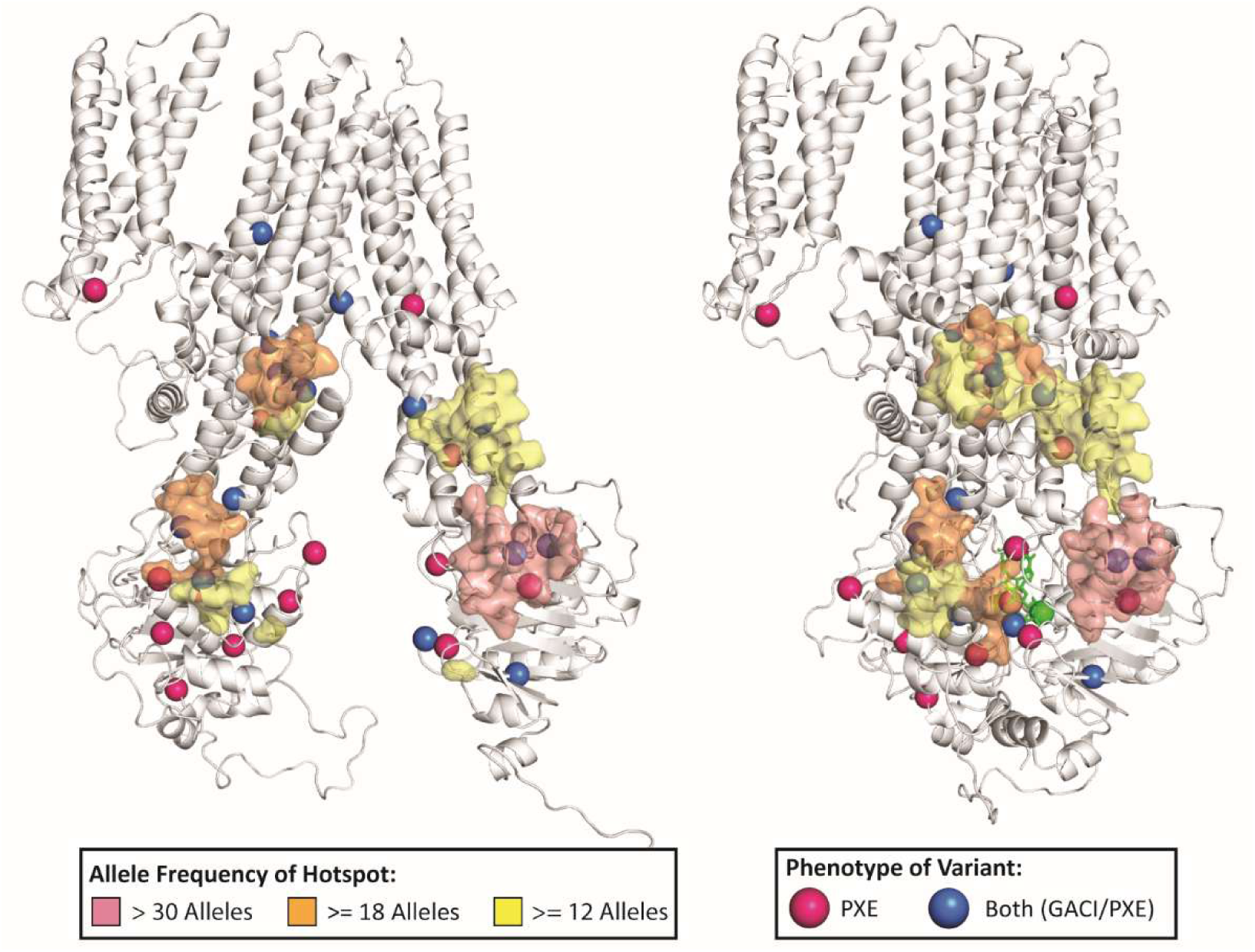
ABCC6 pathogenic variants are highly clustered within three distinct regions. The hotspots of ABCC6 pathogenic alleles, using only allele counts of pathogenic variants on gnomAD, where red areas have more than 30 alleles, orange areas have 17-30 alleles, and yellow regions have 12-16 alleles. Variant locations are marked with a sphere, which is pink when associated with PXE and blue when associated with GACI/PXE. Structures are of State 1 and State 3.

### A Large Portion of Variants of Unknown Significance May Damage ABCC6 Function

We could confidently categorize a large percentage (319 variants; 41% of the total) of VUS by their likely effect on ABCC6’s core functions (**Figure 7**). We can now say how each likely damages ABCC6 function and, therefore, if it is likely to be disease-causing. Further, of the 168 VUS associated with either phenotype, according to ClinVar, 49% were predicted by mechanistic effect. Variants associated only with PXE were classified more frequently (65%), while variants associated with both or neither phenotype were less often predicted (37%). Similar to the pathogenic variants, more VUS were destabilizing (27%) than stabilizing (9%). Further, while most variants within each state have about zero change in structural stability, across all three states, almost all variants have a non-zero structural stability shift. As such the main impact of many variants is on structural stability, especially the PXE variants.

**Figure 7:**
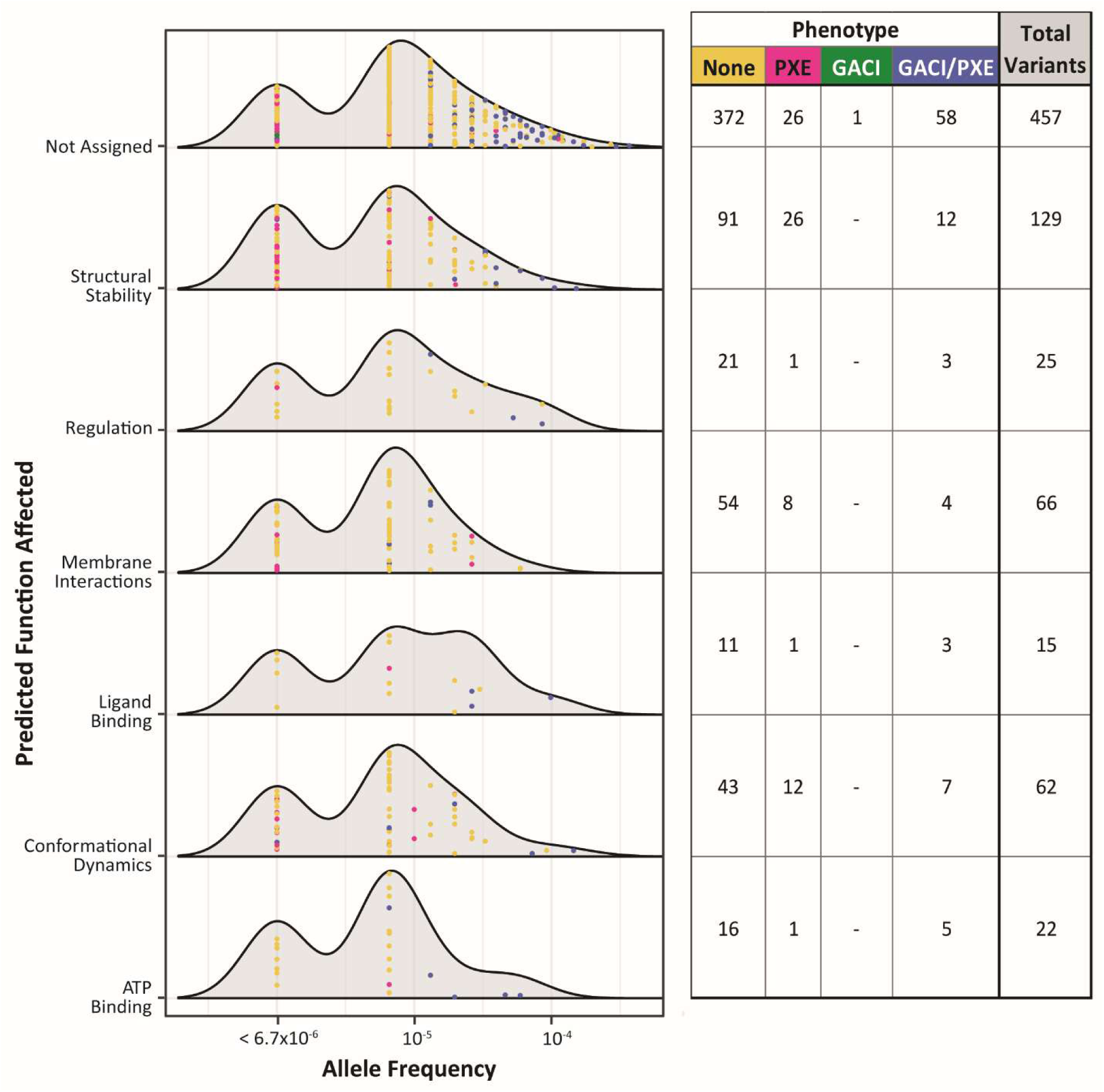
Most VUS were predicted, with most affecting structural stability. Variants are shown across allele frequency and separated by the function of ABCC6 that they likely impact, with the individual variants shown within the distribution colored by the phenotype associated with the variant. Total counts for variants within each effect group are shown on the right.

### A Lack of Clear Differences Between GACI and PXE Variants

Reviewing all 860 distinct non-benign missense variants, only one was associated with GACI alone. For the other variants, 131 were associated with PXE, and another 111 were associated with both GACI and PXE diagnoses (which we will refer to as GACI/PXE variants). Further, for both the pathogenic variants and VUS, the allele frequency was, on average, higher in variants associated with GACI/PXE than those associated with just PXE or neither phenotype **(Figure 8)**. Despite these apparent differences in the clinical severity and allele frequency between the variants responsible for PXE and those for GACI, the lack of specific genetic variants that are unique to GACI makes it challenging to delineate PXE-causing from GACI-causing variants.

**Figure 8:**
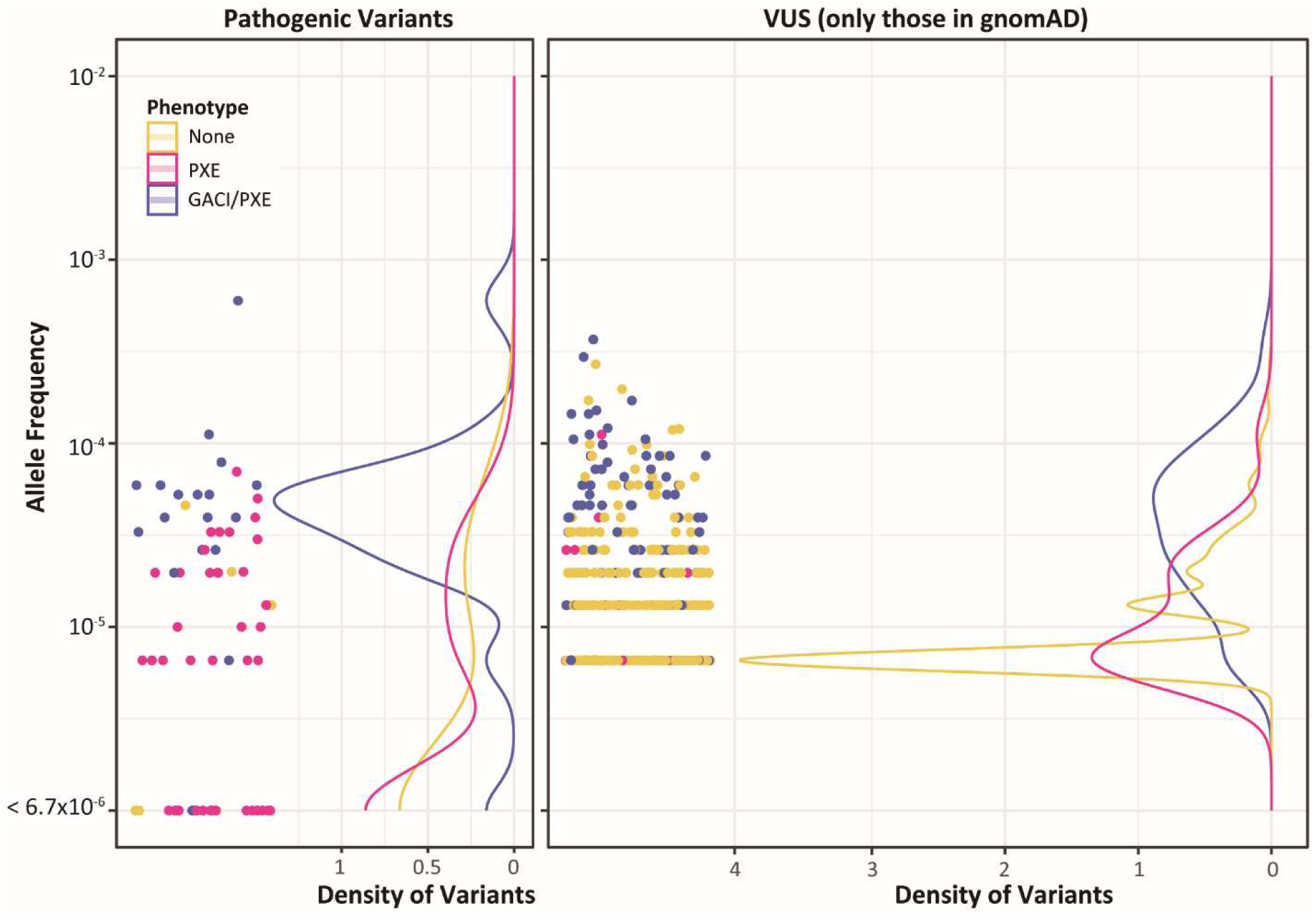
Allele Frequencies for both Pathogenic and Uncertain Significance Variants are higher in variants that cause GACI/PXE than variants that cause PXE alone. On the left, missense pathogenic variants are shown by their allele frequency, whereas variants found only in the ClinVar dataset with no known allele frequency are at the bottom of the plot. The density of variants across the allele frequency range reveals that most GACI/PXE variants are rarer than 1×10^-4^. In contrast, most PXE-only variants have lower allele frequencies or are unobserved in the healthy population. On the right, all missense VUS from the gnomAD dataset are similarly arranged by their allele frequency, revealing once again that the density of PXE variants is greatest at a lower allele frequency than variants that cause GACI/PXE.

### Structural Bioinformatics Links VUS Mechanisms to Pathogenicity

Applying the ACMG guidelines to the variants we studied, despite the lack of segregation, allelic, or familial data, we find 30 VUS that have enough evidence to reclassify as likely pathogenic – a 36% increase in the number of Pathogenic/Likely Pathogenic variants from previously (**Supplementary Text, Supplementary Data**). These proposed reclassifications are due to the novel PM1 or PP3 evidence we provide herein. Furthermore, another 134 VUS will only require one additional line of evidence to be reclassified as likely pathogenic (such as PP1, PP2, PM6, PM2, PM3, or adding new data not available at the time of this work such as new phenotype-genotype associations). This not only expands the number of known likely pathogenic variants clinicians can use to diagnose their patients, but it also enables clinicians to diagnose their patients more efficiently by providing PM1, PM2, PM5, PP3, and PP4 evidence that can be combined with any evidence they collect to create a diagnosis. Furthermore, using the same six levels of evidence, we recommend classifying 50 of the 84 pathogenic (or likely pathogenic) variants in ClinVar as likely pathogenic, showing congruency despite lacking clinical data.

In summary, this study elucidates complex mechanisms underlying ABCC6 dysfunctions and underscores the need for increasing our understanding of population-specific genetic variations that are applicable to n=1 cases. We conclude that integrating biophysical and biochemical metrics, proteogenomic approaches, and computational algorithms has substantially enhanced our comprehension of ABCC6 variants. This understanding builds a trajectory for more accurate and mechanistic clinical diagnoses, leaving less ambiguity for patients.

## Discussion

Currently, less than 14% of *ABCC6* alleles reported in ClinVar and gnomAD have a confident clinical interpretation. Among variants with an interpretation, there is no clear underlying disease mechanism. These gaps in knowledge make variant interpretation and treatment difficult. To further investigate this issue, we used a multi-tiered proteogenomic approach to study ABCC6 variants. We analyzed the variants in three steps to characterize their molecular context and mechanistic effect. Our findings show that our approach is effective and insightful in determining the functional mechanism of ABCC6 genetic variants. By encoding this process as an algorithm, we mechanistically characterized 41% of VUS to high confidence (See **Supplementary Materials, Table S1**). Further, with the information gathered, we also explored differences between variants associated with GACI and PXE - differences that may help us better understand the relationship between the two fundamentally similar yet distinct disorders. Projecting currently identified pathogenic variants onto our 3D structures enabled us to find mutational hotspots. Finally, we provide evidence for five ACMG categories across all 901 variants based on novel structural biology insights, which can be used to aid in classification. Our study adds to what is currently known about both the mechanistic effect of ABCC6 variants and the deleteriousness of VUS, which can help clinicians better interpret patient genotypes.

Previous studies of ABCC6 variants have relied on a singular homology model, characterized up to tens of variants, or have relied on purely functional information. This is the first study of ABCC6 to describe nearly a thousand variants using multi-state models and extensive mechanistic data. This integrative approach enables us to characterize far more variants than we could if we used one model or only functional information. Our results demonstrate the utility of our multi-tiered proteogenomic approach and lead to novel conclusions about ABCC6.

Few studies have directly compared PXE-causing variants to GACI-causing variants in ABCC6; therefore, our study boasts new information about ABCC6 variants’ disease associations. We discovered few variants observed exclusively in GACI patients and few differences in mechanisms between variants associated with PXE and those associated with GACI/PXE. This may, in part, be due to lack of specificity in ClinVar variant reporting, especially when a variant is added by large-scale genetics companies which don’t provide phenotypic information. Further, we found that GACI/PXE variants have a higher allele frequency. Since GACI is less common than PXE, GACI would be more statistically likely to be associated with more common variants if all variants were equally likely to lead to GACI on their own^57, 58^. Altogether, this suggests that the molecular differences between PXE and GACI are not caused by specific variants, and other distinguishing factors are likely at play, such as a genetic or epigenetic modifier, a metabolic environment, a change in regulation, an external stressor, etc. This directs future studies of ABCC6 to focus on patient-to-patient differences, rather than variant-to-variant differences, in differentiating GACI and PXE.

This study also represents a significant advance in understanding genetic variants in *ABCC6* through methodologies that could help with other rare disease genes. Our approach predicted the mechanistic effect of 319 VUS that likely impair ABCC6 function enough to cause protein dysfunction. While our accuracy and precision are more balanced compared to current best-in-class pathogenicity prediction algorithms, the mechanistic nature of our structure-based classification informs potential treatment options. It enables hypothesis testing, which improves over standard genomic *in silico* scores.

Another significant contribution of this study is its methodology, which can be applied to many other proteins and variants. Indeed, our study takes a new approach to genomic variant analysis and interpretation, focusing on the mechanistic effect of a variant on the translated gene product rather than the likelihood of pathogenicity. The damaging potential of a variant is clearly related to its probability of pathogenicity. However, our current calculations do not capture all contexts (such as changes to protein folding or impacts on any potential protein interactions that have yet to be discovered). Creating an algorithm specifically for ABCC6 that considers a large amount of specific contextual information enables even the rare and singleton variants to be understood – something statistics-based approaches struggle with. Furthermore, our study revealed clusters of pathogenic variants within three main hotspots, which are spatially near functionally significant zones of ATP binding and ligand binding. This demonstrates the close relationship between a variant’s pathogenicity and the protein’s function – making understanding protein function especially critical in advancing better variant interpretation. Our hotspot analysis and functional annotations enabled much of our variant reclassification and will continue to provide PM1 evidence for further reclassification of VUS. Overall, our results demonstrate that a protein-specific structural bioinformatics approach like ours helps us better understand the mechanisms of pathogenicity for VUS, and this methodology can be used to study variants of any protein with some known structural and functional information.

Through future studies, additional cell-based data will further understanding of the differences between PXE and GACI. Using multiple functional conformations to structurally characterize variants, integrated with additional functional genomics information, can create more specific and accurate studies. Results from the current work can guide future studies of other variants with the methods described herein, leading to better interpretation of patient genomes and, eventually, more successful treatment of both diseases.

## Conclusion

With a large range of phenotypes emerging from variants in *ABCC6*, determining the deleteriousness of patient variants and their mechanisms for impairing function is critical for diagnostic and therapeutic research. The current study employs a novel multi-tiered approach to ABCC6 variant analysis and identifies the underlying molecular effects for 60% of pathogenic variants and 43% of VUS, identifying additional information that supports 30 VUS to be reclassified as Likely Pathogenic. Approaches such as we report here not only enable more genetic diagnoses, but also provide a new kind of information for clinical interpretation that is applicable to the one-of-a-kind genetic variation that frequently underlies rare diseases. Finally, most variants were predicted to impact the structural stability of ABCC6, highlighting the importance of considering the structural implications of genetic variants on their encoded products. This study supports a different way of thinking about variant interpretation in general: the mechanistic effects of DNA changes can be enhanced by closely considering the intersection of structure and function.

## Data Availability

All data used in assigning variant effects and the assigned variant effect and pathogenicity is available in the supplementary table. In the supplementary text, we also include the code for the algorithm used to make our structure-based predictions and our derived ACMG-based classifications. Models of ABCC6 in three states are on ModelArchive.

## Acknowledgments

This research was completed in part with computational resources and technical support provided by the Research Computing Center at the Medical College of Wisconsin.

## Author Contributions

Conceptualization: W.G., S.Z., R.U., M.Z., N.H., B.R.; Data curation: S.Z., J.W.; Formal analysis: J.W., S.J.; Investigation: J.W.; Methodology: J.W., N.H., S.J., B.R., S.Z., M.Z.; Software: J.W., N.H., B.R., M.Z.; Supervision: W.G., R.U., M.Z.; Visualization: J.W., N.H., M.Z.; Writing-original draft: J.W.; Writing-review & editing: J.W., N.H., B.R., M.Z.

