## Supplemental Text for "A Multi-State Structural Genomics Approach Enables Large-Scale, Mechanistic, and Context-Specific Classification of ABCC6 Genetic Variants Implicated in Calcification Diseases"

### Supplementary Materials

Table S1: **Information, scores, and ACMG evidence and classification for all variants.** Supplementary_Table_S1.xlsx (see other files). In this file, each variant analyzed is listed, along with all data used in analysis and our results. Below is a description of each column.

1. HGVS cDNA: the HGVS description of the variant at the cDNA level.
2. HGVS protein: the HGVS description of the variant at the amino acid level.
3. Allele frequency: the global allele frequency for the variant, as per gnomAD v3.
4. Clinical significance: the current variant interpretation, as per ClinVar. For simplicity’s sake, variants which were labeled as pathogenic, likely pathogenic, or pathogenic/likely pathogenic were all simplified to “Pathogenic/Likely Pathogenic”; the same was done with benign and likely benign variants.
5. Phenotype: the simplified associated phenotype, as per ClinVar. All phenotype descriptions which include pseudoxanthoma elasticum (including more informative descriptions such as “autosomal recessive inherited pseudoxanthoma elasticum”, etc.) were simplified to “PXE”, descriptions including generalized arterial calcification of infancy were simplified to “GACI”, and descriptions with both were simplified to “Both”.
6. State 1 Stabilization: FoldX’s change in free energy of folding (in kcal/mol) for the State 1 structure of ABCC6 for each variant, in comparison to the WT State 1 structure.
7. State 2 Stabilization: FoldX’s change in free energy of folding (in kcal/mol) for the State 2 structure of ABCC6 for each variant, in comparison to the WT State 2 structure.
8. State 3 Stabilization: FoldX’s change in free energy of folding (in kcal/mol) for the State 3 structure of ABCC6 for each variant, in comparison to the WT State 3 structure.
9. Number of Damaged States: a string describing how many states are destabilized or stabilized (having a change in free energy of folding of > 1.8 or < -1.8).
10. PAM: the amino acid change’s score from the Point accepted mutation (PAM) matrix.
11. Hydrophobicity Change (Kyte-Doolittle): the difference in Kyte-Doolittle hydrophobicity score between the original and alternative amino acid.
12. Location-Based Annotations: all functional information about residues within 5 residues and/or within XX Å of the variant location.
13. Domain	: which functional domain the protein family is in.
14. Predicted Affected Function: which mechanism of ABCC6 WT functioning we predict the variant may be impacting, if any.
15. PM1: whether this variant has the ACMG PM1 criterion for being pathogenic or not.
16. PM2: whether this variant has the ACMG PM2 criterion for being pathogenic or not.
17. PM5: whether this variant has the ACMG PM5 criterion for being pathogenic or not.
18. PP2: whether this variant has the ACMG PP2 criterion for being pathogenic or not.
19. PP3: whether this variant has the ACMG PP3 criterion for being pathogenic or not.
20. PP4: whether this variant has the ACMG PP4 criterion for being pathogenic or not.
21. Suggested Clinical Significance: the existing clinical significance (column 4) was updated such that when a variant has enough evidence to be reclassified as “Likely Pathogenic”, it is now labeled as so. When a variant has almost enough evidence to be reclassified as likely pathogenic, its previous classification is annotated with a star.

Code: Algorithm for Variant Effect Categorization

This code takes, as input, the first 12 columns from the supplementary table as that is all the data we used in assigning variant effect. If other variants were to be analyzed, the same 12 columns of information would need to be provided with the given names (but not necessarily the same order).

```
df <- read_excel("Supplementary_Variant_Data_And_Results_New.xlsx")[, c(1:13)]
colnames(df) <- c("HGVSc", "HGVSp", "AF2", "Category", "Pheno", "State1", "State2", "State3",
                  "damagedState", "PAM", "HydrophobicityChange", "Annotation", "Location")

df <- df %>% rowwise() %>%
  mutate(Alt = str_sub(HGVSp, -1)) %>%
  mutate(Effect =
         if(grepl("ATP-adjacent", Annotation, fixed = TRUE) &
            (State1 >= 1.5 | State2 >= 1.5 | State3 >= 1.5 | PAM > 3 |
             PAM < -2 | HydrophobicityChange > 2 | HydrophobicityChange < -2 )) {"ATPbinding"}
         else if(grepl("ATP-mot", Annotation, fixed = TRUE) & (PAM > 2 | PAM < -2)){"ATPbinding"}
         else {NA}) %>%
  mutate(Effect =
         if(is.na(Effect)){
             if(grepl("Ligand", Annotation, fixed=TRUE) & (PAM > 2 | PAM < -1)){"Ligandbinding"}
             else{NA} }
         else{Effect}) %>%
  mutate(Effect =
         if(is.na(Effect)){
             if(grepl("Membrane", Annotation, fixed=TRUE) &
                ( (HydrophobicityChange > 0.5) | (Alt == "R" | Alt == "K") |
                  ( (State1 >= 1.8 | State2 >= 1.8 | State3 >= 1.8) & damagedState == "Multiple") ) )
               {"MembraneInteractions"}
             else{NA} }
         else{Effect}) %>%
  mutate(Effect =
         if(is.na(Effect)){
             if ( (grepl("Phosphorylation", Annotation, fixed = TRUE) |
                   grepl("Acetylation", Annotation, fixed = TRUE) |
                   grepl("Ubiquitination", Annotation, fixed = TRUE) |
                   grepl("Regulatory", Annotation, fixed = TRUE) )
                  & ( PAM < -2 | PAM > 3) ) {"Regulation"}
             else{NA} }
         else{Effect}) %>%
  mutate(Effect =
         if(is.na(Effect)){
           if (State1 >= 1.8 & damagedState == "One" & startsWith(Location, "TMD") |
               State2 >= 1.8 & damagedState == "One" & startsWith(Location, "TMD") |
               State3 >= 1.8 & damagedState == "One" & startsWith(Location, "TMD") |
               State1 <= -1.8 & damagedState == "One" & startsWith(Location, "TMD") |
               State2 <= -1.8 & damagedState == "One" & startsWith(Location, "TMD") |
               State3 <= -1.8 & damagedState == "One" & startsWith(Location, "TMD") ) {"Conformation"}
           else{NA} }
         else{Effect}) %>%
  mutate(Effect =
         if(is.na(Effect)){
             if (State1 >= 1.8 | State2 >= 1.8 | State3 >= 1.8) {"Stability"}
             else if (grepl("NBD-TMD_joint_instability", Annotation, fixed=TRUE) &
                      (State1 >= 1 | State2 >= 1 | State3 >= 1)){"Stability"}
             else if (grepl("NBD-instability", Annotation, fixed=TRUE) &
                      (State1 >= 1 | State2 >= 1 | State3 >= 1)){"Stability"}
             else{NA} }
         else{Effect})
```

Text:

The 3D hotspot consists of the following residues: 394, 483, 484, 503, 513, 514, 515, 516, 517, 521, 693, 695, 696, 761, 762, 763, 764, 766, 781, 1032, 1109, 1110, 1111, 1112, 1113, 1115, 1116, 1117, 1118, 1119, 1133, 1134, 1135, 1136, 1137, 1143, 1160, 1161, 1272, 1299, 1307, 1308, 1309, 1310, 1311, 1312, 1313, 1315, 1316, 1317, 1318, 1330, 1331, 1335, 1338, 1339, 679, 694, 776, 780, 231, 232, 233, 486, 502, 711, 726, 754, 755, 756, 757, 758, 783, 926, 1228, 1231, 1232, 1235, 1246, 1323, 700, 808.

Previously unclassified variants that we suggest be classified as Likely Pathogenic include: A1303P, A1318G, A766D, D718G, D777N, E1400K, G1133A, G1133C, G1296D, G1299S, G1302R, G1321S, G1354R, G1405S, G1501S, G663C, G666W, G755R, K502R, L1335P, L1335Q, L355R, L495H, L673P, L677P, L726P, M751K, Q1347H, Q1406K, Q698P, R1114C, R1114H, R1114P, R1138P, R1138Q, R1235Q, R1235W, R1314W, R1339C, R1339H, R600C, R760W, R765Q, R765W, R807W, S1307P, T1301I, T811M, T811R, V1298F, P664S, A1303T, A1318T, A766V, E521D, E699G, F1493L, G1299R, G1311E, G1481S, G1501C, G663R, G663S, I1424T, K502M, L1063P, L753P, P1346S, P1483L, P1483Q, Q1406H, R1235G, R1339L, R1339S, R1459C, R600L, R807G, S754C, T811A, V1404M, V810M.
